# Nitrapyrin has far reaching effects on the soil microbial community structure, composition, diversity and functions

**DOI:** 10.1101/2020.07.21.205765

**Authors:** Ruth Schmidt, Xiao-Bo Wang, Paolina Garbeva, Étienne Yergeau

## Abstract

Nitrapyrin is one of the most common nitrification inhibitors that are used to retain N in the ammonia form in soil to improve crop yields and quality. Whereas the inhibitory effect of nitrapyrin is supposedly specific to ammonia oxidizers, in view of the keystone role of this group in soils, nitrapyrin could have far-reaching impacts. Here, we tested the hypothesis that nitrapyrin leads to large shifts in soil microbial community structure, composition, diversity and functions, beyond its effect on ammonia-oxidizers. To test this hypothesis, we set-up a field experiment where wheat (*Triticum aestivum* cv. AC Walton) was fertilized with ammonium nitrate (NH_4_NO_3_) and supplemented or not with nitrapyrin. Rhizosphere and bulk soils were sampled twice, DNA was extracted, the 16S rRNA gene and ITS region were amplified and sequenced to follow shifts in archaeal, bacterial and fungal community structure, composition and diversity. To assess microbial functions, several genes involved in the nitrogen cycle were quantified by real-time qPCR and volatile organic compounds (VOCs) were trapped in the rhizosphere at the moment of sampling. As expected, sampling date and plant compartment had overwhelming effects on the microbial communities. However, within these strong effects, we found statistically significant effects of nitrapyrin on the relative abundance of Thaumarchaeota, Proteobacteria, Nitrospirae and Basidiomycota, and on several genera. Nitrapyrin also significantly affected bacterial and fungal community structure, and the abundance of all the N-cycle gene tested, but always in interaction with sampling date. In contrast, nitrapyrin had no significant effect on the emission of VOCs, where only sampling date significantly influenced the profiles observed. Our results point out far-reaching effects of nitrapyrin on soil and plant associated microbial communities, well beyond its predicted direct effect on ammonia-oxidizers. In the longer term, these shifts might counteract the positive effect of nitrapyrin on crop nutrition and greenhouse gas emissions.

## Introduction

Together with phosphorus, nitrogen (N) is the most limiting nutrient for plant growth, especially in agroecosystems. One of the key advances of the green revolution was the intensive use of inorganic fertilizers to increase crop yields. However, fertilization, especially with N, has become at the center of the environmental and sustainability issues face by agriculture. The adverse effects of the indiscriminate use of N fertilizers range from N losses through ammonia volatilization, nitrate leaching and denitrification, leading to atmospheric and groundwater pollution by nitrous oxide and nitrate (1). A lot of these adverse effects are linked to soil microbial processes such as nitrification and denitrification.

Nitrification is the oxidation of ammonia to nitrate carried out by chemoautotrophs that use the energy thereby generated to fix atmospheric CO_2_. Ammonia oxidation, the first and rate-limiting step of nitrification that converts ammonia to nitrite (2, 3) is carried out by ammonia oxidizing bacteria (AOB), belonging to the genera *Nitrosomonas*, *Nitrosospira,* and *Nitrosococcus* and the more recently discovered ammonia oxidizing archaea (AOA) of the Thaumarchaeota phylum (4) belonging to the genera including *Nitrososphaera*, *Nitrosopumilus*, *Nitrosoarchaeum* and *Cenarchaeum* (5). These keystone species are ubiquitously found in soil (6), with AOA generally being more abundant than AOB (7, 8). An increasing number of evidence has shown that AOA may play a more important role than AOB in nitrification (9, 10). Yet, the relative contribution of AOA and AOB to nitrification rates remain unclear, largely depending on soil conditions, including pH (10), temperature (11) and N availability (12).

However, the transformation by nitrifiers of ammonia to nitrate to generate energy is environmentally and agriculturally unfavorable. Nitrate, because of its negative charge, is more mobile in soil and more prone to leaching than ammonia, and at the same time nitrate is the substrate for denitrification, which, if incomplete, can lead to the emission of the potent greenhouse gas nitrous oxide. Nitrate, although more mobile, is energetically less favorable for plant growth, as it needs to be actively taken up, and transformed back to ammonia for protein synthesis (13). This is especially important for cereals, where grain quality is often linked to its protein content. In order to counteract the plant and environmental adverse effects of nitrification, nitrification inhibitors (NI) have been used in recent decades in combination with ammonia-based fertilizers. Out of these, nitrapyrin (2-chloro-6-(trichloromethyl)-pyridine) is one of the most commonly applied commercial NIs and was shown to increase soil N retention (14) and crop yields in a variety of crops (15), while decreasing N_2_O emissions (16). On average, crop yields were shown to increase by 7% and soil N retention by 28%, while N leaching decreased by 16% and greenhouse gas emissions decreased by 51% (15). A meta-analysis of globally representative NI research trials found that nitrapyrin increased grain production by 2% - 9% for wheat and maize, respectively (17).

Nitrapyrin delays nitrification by temporarily deactivating the ammonia monooxygenase (AMO), the enzyme responsible for ammonia oxidation (18, 19). The α subunit of AMO is encoded by the *amoA* gene, which is homologous in AOA and AOB (20), suggesting that both AOA and AOB could be inhibited by nitrapyrin. However, studies investigating the inhibitory effects of nitrapyrin on AOA and AOB (determined by *amoA* gene abundance) have found varying results. For instance, AOB were reported to be not affected by nitrapyrin, despite a decrease in nitrification rates (21). In another study, an inhibition of AOA growth was suggested by a lack of increase in the abundance of the *amoA* gene (Lehtovirta-Morley *et al.*, 2013). Similarly, Shen *et al.* (2013) found a very weak inhibitory effect of nitrapyrin on the AOB, but a moderate effect on the AOA. Interestingly, the inhibition of AOA by nitrapyrin was shown to lead to a reduction in the AOA to AOB ratio in soils with low nitrogen use efficiency, which was positively correlated with potential nitrification rates (24). Overall, these studies suggest that nitrapyrin shifts the AOA to AOB ratio, thus inhibiting nitrification at different rates depending on the relative contribution of AOA and AOB to nitrification. This shift is probably linked to environmental and soil physico-chemical characteristics. Yet, it remains unresolved whether a shift in these keystone species has a cascading effect on the overall microbial community structure, composition, diversity and functions, and how this varies through the growing season.

Soil microbes, including nitrifiers, produce a plethora of volatile organic compounds (VOCs) that are small carbon-based molecules with high vapor pressure and low boiling point (25, 26). Due to these unique characteristics, this interesting group of metabolites mediates microbial below-ground interactions (27–29) and fulfills important functions in terrestrial ecosystems (30, 31). The production of VOCs is strongly affected shifts in the community composition (32), and is even sensitive to small-scale changes in low abundant microbes (33). Yet, how nitrapyrin affects microbial VOC emission, either directly or indirectly has not been investigated.

Therefore, we hypothesized that nitrapyrin shifts the abundance of the overall microbial community structure, composition, diversity and functions. The objectives of this study were to explore the effects of nitrapyrin applied on field-grown wheat on 1) the overall microbial community structure, composition and diversity, 2) the abundance of genes encoding for enzymes involved in ammonia oxidation, N-fixation, ammonia oxidation and denitrification, and 3) the rhizosphere VOC emissions.

## Materials and methods

### Field experiment and sampling

The field trial was set up in June 2019 at the Armand-Frappier Santé Biotechnologie Research Center (Laval, Québec, Canada) (Figure S1). A total of twelve plots (1 m × 1 m) were established based on a randomized design with six replicated plots per treatment. Approximately 20 g of wheat seeds (*Triticum aestivum* cv. AC Walton) were sown in 4 rows in each plot (Figure S1). A solution of ammonium nitrate (NH_4_NO_3_) and the nitrification inhibitor nitrapyrin,2-Chloro-6-pyridine (NI) were applied twice during the growing season on the soil surface. The first time (June 11^th^, 2019) 10 g N (28.57g ammonia nitrate) was applied for each plot and 0.05 g NI for each NI-treated plot. The second time (July 10^th^, 2019), 10 g N was applied for each plot and 0.1 g NI for each NI-treated plot. Weeds were removed every week. Sampling of bulk soil and rhizosphere soil, as well as entrapment of VOCs using PDMS tubes (see below for more details) in the rhizosphere was conducted twice, at the stage of grain filling (July 23^rd^, 2019) and at the time of harvest (September 5^th^, 2019), totalling 48 samples for DNA extraction (2 treatments × 2 compartments × 2 sampling dates × 6 replicates = 48 samples) and 24 samples for VOC analyses (2 treatments × 1 compartment × 2 sampling date × 6 replicates = 24 samples). For bulk soil, five 2-cm diameter soil cores from the upper 5-cm were collected from each plot and pooled together to create one composite sample. Rhizosphere soil was collected by shaking off soil from one plant in the center of each plot. Soils were sieved (2.0 mm mesh size) and placed into a sterile plastic bag and immediately stored at −80°C prior to DNA extraction. VOCs were collected with Rotilabo®-silicone tubes (PDMS tubes; Carl Roth GmbH+Co. KG, Karlsruhe, Germany), which were placed in the rhizosphere of one uprooted plant per plot (Figure S1). The PDMS tubes were pre-treated as described previously (34). After 20 minutes, PDMS tubes were removed and kept at −20□°C until analysis.

### DNA extraction and PCR amplicon sequencing

Soil DNA was extracted from 0.3 g of well-mixed soil for each sample using the DNeasy PowerLyzer PowerSoil Kit (Qiagen), following the manufacturer’s instructions. Extracted genomic DNA was quantified using a PicoGreen™ (Thermo Scientific, OR, U.S.A.) assay (Ahn, Costa et al. 1996) with a Tecan Infinite M1000 PRO (Thermo Scientific). DNA was stored at −20°C until used for PCR.

PCR amplicon libraries were prepared for the archaea and bacterial 16S rRNA gene using primers 515F and 806R targeting the V4 region (35) and for the fungal ITS1 region using primers ITS1F and 58A2R (36). The first step of PCR contained the template specific primers with a short adaptor sequence, and the second step of PCR was conducted with primers containing the Illumina barcodes. Two PCR amplification steps were performed in a T100™ Thermal Cycler (Bio-Rad, U.S.A.). Reagents and amplification conditions for each of the PCR are shown in Table S1. Amplicons were verified on 1% agarose gels and purified using AMPPure XP beads (Beckman Coulter, Indianapolis, U.S.A.) following the manufacturer’s instructions. Sequencing was conducted on an Illumina MiSeq sequencer (2 × 250 pair-end) at the McGill University and Genome Québec Innovation Center (Montréal, Canada). A total of 7,416,206 16S rRNA gene reads and 6,649,643 ITS region reads were produced. The raw datasets and associated metadata are available through NCBI BioProject accession PRJNA634744.

### Quantitative real-time PCR (qPCR)

qPCR was performed using the iTaq universal SYBRGreen^®^ kit following the manufacturer protocol (Bio-Rad Laboratories Inc, Hercules, CA) on an Agilent Mx3005P qPCR Systems (Agilent Technologies, CA, U.S.A.). Primers crenamoA23f and crenamoA616r (37), *amoA*1f and *amoA*2r (38), Cd3af and R3cd (39), *nirK*876f and *nirK*1040r (40) and nifHPo1f and nifHPo1r (41) were used for quantification of the archaea ammonia monooxygenase A subunit gene (*amoA*-AOA hereafter), bacterial ammonia monooxygenase A subunit gene (*amoA*-AOB hereafter), *nirS*, *nirK* and *nifH* genes, respectively. Standards were prepared prior to qPCR for selected functional genes as described previously (42). Briefly, the genes of interest were amplified from DNA extracted from an agricultural soil, and the resulting amplicons were purified and concentrated using a Wizard SV gel purification kit (Promega Corporation, U.S.A.), and cloned into a P-Gem T plasmid (Promega Corporation, U.S.A.). The plasmids were linearized with the restriction enzyme SacII, quantified using a fluorescent DNA-binding dye (Quanti-iT™PicoGreen™ ds Assay, Life Technologies, U.S.A.) and then serially diluted (10^1^–10^8^ copies μl^−1^) for creating a standard curve. Reagents and reaction conditions for qPCR are shown in Table S2.

### VOC analysis

VOCs were desorbed from the PDMS tubes by using an automated thermodesorption unit (model UnityTD-100, Markes International Ltd., UK) at 250°C for 12 min (He flow 50 ml/min). The desorbed VOCs were subsequently collected on a cold trap at −10°C and introduced into a GC-QTOF (model Agilent 7890B GC and the Agilent 7200AB QTOF, USA) by heating the cold trap for 10 min to 280°C. A split ratio was set to 1:10. The column used was a 30 × 0.25 mm IDDB-5MS with as film thickness of 0.25 μm (Agilent 122-5532, USA). The temperature program was as follows: 2 min at 39°C, 3.5°C/min to 95°C, 4°C/min to 165°C and finally 15°C/min to 280°C that was hold for 15 min. VOCs were detected by the MS operating at 70 eV in EI mode. Mass spectra were acquired in full scan mode (30–400 AMU, 4 spectras/sec). Collected GC/MS data was converted to mzData files using the Chemstation B.06.00 (Agilent Technologies, Santa Clara, USA) and further processed (peak picking, baseline correction and peak alignment) in an untargeted manner with MZmine 2 (43). Detected compounds were identified using NIST-MS Search by comparing the spectra, accurate mass, linear retention indices and spectra match factor with NIST 2014 V2.20 (National Institute of Standards and Technology, USA, http://www.nist.gov), Wiley 9th edition, and in-house spectral libraries. Putative compounds were identified using AMDIS 2.72 (National Institute of Standards and Technology, Gaithersburg, USA) and linear retention indices of VOCs were calculated according to the method described by Strehmel *et al.*, 2008. PDMS contaminants were removed from the final list.

### Bioinformatics analysis

Primers were trimmed with up to one mismatch allowed and starting position ≤1. Forward and reverse reads of same sequence were merged with at least 30 bp overlap and <0.25 mismatches by using FLASH v1.2.5 (45). The sequences were then quality trimmed using Btrim (46) with threshold of QC > 30 over 5bp window size. Merged sequences with an ambiguous base, or < 240 bp for 16S and < 200 bp for ITS were discarded. Chimera sequences were detected and removed with the UCHIME algorithm (47). Sequences were clustered into operational taxonomic units (OTUs) with a 97% identity cut-off using UPARSE (48) and their taxonomic affiliation was assigned using RDP 16S rRNA reference database (49) and UNITE ITS reference database (50). The above mentioned steps were performed using an in-house pipeline that was built on the Galaxy platform at the Research Center for Eco-Environmental Sciences, Chinese Academy of Sciences (http://mem.rcees.ac.cn:8080/) (51).

### Statistical analysis of 16S and ITS data

All statistical analysis and figure generation were performed in RStudio v1.2.5042 running R v3.6.3. 16S and ITS data were analysed using the phyloseq package v 1.28.0 (52), the microbiome package v 1.6.0 (53) and the microbiomeutilities package v 0.99.00 (54). Shannon diversity index, Inverse Simpson index matrices were calculated with the *estimate_richness* function of the phyloseq package v 1.28.0. Faith’s Phylogenetic Diversity (PD) (55) was estimated using the *pd* function of the picante package v 1.8.1 (56). Three-way repeated measures analysis of variance (ANOVA) was performed for alpha diversity indices across treatment, sampling date and compartment using the *aov* function of the stats package v 3.6.3 (57). After removal of singletons and unclassified phyla, top 10 phyla and genera were extracted using the *taxa_sums* function of the phyloseq package v 1.28.0. Three-way repeated measures ANOVA was performed for the top 10 phyla and genera across treatment, sampling date and compartment using the *aov* function of the stats package v 3.6.3. Principal Coordinates Analysis (PCoA) with Bray-Curtis dissimilarity was used to visualize similarity between samples based on bacterial and fungal composition using the *ordinate* function of the phyloseq package v 1.28.0. The effects of the treatment, sampling date and compartment on the bacterial and fungal community structure were tested using permutational multivariate analysis of variance (PERMANOVA) with the *adonis* function based on Bray–Curtis dissimilarity indices of the vegan package v 2.5-6 (58). All plots were generated using the ggplot2 v 3.3.0 (59).

### Statistical analysis of qPCR data

Values (log copies per g soil) were first tested for homogeneity of variance using the *leveneTest* function of the car package v 3.0-7 (60), followed by Shapiro-Wilk’s normality test using the *shapiro_test* function of the rstatix package v 0.4.0 (61). The non-parametric *krusk.test* function of the of the stats package v 3.6.3 (57) was used to compare samples, followed by posthoc analysis using the *dunnTest* function of the FSA package v 0.8.30 (62). All plots were generated using the ggplot2 v 3.3.0 (59)

### Statistical analysis of VOCs

Aligned m/z features were filtered and normalised as time-series data using the metaboAnalystR package v 2.0.1 (63) as follows: features containing a 0.5% cut-off of missing values were removed, missing values with a minimum positive value (here: 2463.451875). Further feature filtering was performed based on Interquantile Range. Finally, data were normalised by median, log transformed and auto scaled. The normalised data were then used to perform ANOVA (*P* < 0.01) and Principal Component Analysis (PCA) using the metaboAnalystR package v 2.0. The effects of treatment and sampling date as well as their interaction effect on the overall VOC composition were tested using PERMANOVA with the *adonis* function of the vegan package v 2.5-6 (58) based on Euclidean distances. A heatmap of significant features based on ANOVA was created using the *ggheatmap* function of the heatmaply package 1.1.0 (64) by hierarchical clustering based on Euclidean distances. Values of individual compounds were subsetted and plotted as log median values. All plots were generated using the ggplot2 package v 3.3.0 (59).

## Results

### Diversity, composition and structure of soil microbial communities

The sequences obtained from the 16S rRNA gene and the ITS region were grouped into 26,979 and 4,482 OTUs with a 97% sequence similarity threshold, respectively. Bacterial and archaeal, as well as fungal alpha diversity estimated by the Shannon, inverse Simpson and phylogenetic diversity (PD) indices did not differ significantly between the NI treated samples and the controls (*P* > 0.05) (Table 1). For bacterial and archaeal communities, sampling date significantly affected the Inverse Simpson index (*P* < 0.01) with an increased diversity at the second sampling date (Table 1, Figure1). Compartment significantly affected the Faith’s PD (*P* < 0.01) (Table 1), with a lower diversity in bulk soil as compared to the rhizosphere for the first sampling date for both the NI and control treatments (Figure 1). For fungal communities, compartment significantly affected the Shannon diversity index (*P* < 0.01) and the Inverse Simpson index (*P* < 0.001) (Table 1) with a lower diversity in rhizosphere as compared to the bulk soil (Figure 1). Sampling date also significantly affected the Shannon (*P* < 0.001), the inverse Simpson (*P* < 0.001) and Faith’s PD (*P* < 0.001) indices for fungal communities (Table 1), with a decreased diversity at the second sampling date (Figure 1).

**Table 1:** Three-way repeated measures ANOVA of the bacterial, archaeal (16S) and fungal (ITS) alpha diversity examined by Shannon index, Inverse Simpson and Faith’s PD.

|  | 16S |  |  |  |  |  | ITS |  |  |  |  |  |
| --- | --- | --- | --- | --- | --- | --- | --- | --- | --- | --- | --- | --- |
|  | Shannon |  | Inverse Simpson |  | PD |  | Shannon |  | Inverse Simpson |  | PD |  |
|  | F | P | F | P | F | P | F | P | F | P | F | P |
| Treatment | 0.001 | 0.971 | 0.555 | 0.461 | 1.741 | 0.195 | 0.588 | 0.448 | 0.658 | 0.422 | 0.924 | 0.342 |
| Compartment | 0.281 | 0.599 | 0.612 | 0.439 | 7.977 | **0.007** | 9.523 | **0.004** | 17.086 | < **0.001** | 0.308 | 0.582 |
| Date | 1.175 | 0.285 | 0.016 | **0.003** | 0.378 | 0.542 | 49.577 | < **0.001** | 53.389 | < **0.001** | 26.062 | < **0.001** |
| Treatment × Compartment | 0.02 | 0.888 | 2.838 | 0.1 | 0.121 | 0.73 | 1.314 | 0.258 | 2.237 | 0.143 | 0.74 | 0.395 |
| Treatment × Date | 0.978 | 0.329 | 0 | 0.997 | 0.923 | 0.342 | 1.492 | 0.229 | 0.905 | 0.347 | 0.21 | 0.649 |
| Compartment × Date | 1.552 | 0.22 | 1.273 | 0.266 | 2.485 | 0.123 | 0 | 0.989 | 2.586 | 0.116 | 1.434 | 0.238 |
| Treatment × Compartment × Date | 0.082 | 0.777 | 1.714 | 0.198 | 0.345 | 0.560 | 0.119 | 0.732 | 1.948 | 0.171 | 0.416 | 0.522 |

Treatment: treatment with nitrapyrin and control without nitrapyrin. C: Compartment (Bulk soil and Rhizosphere). Date: sampling dates (2019-08-20, 2019-09-05). *P*-values marked in bold indicate statistically significant difference with $P < 0.05$ .

**Figure 1:**
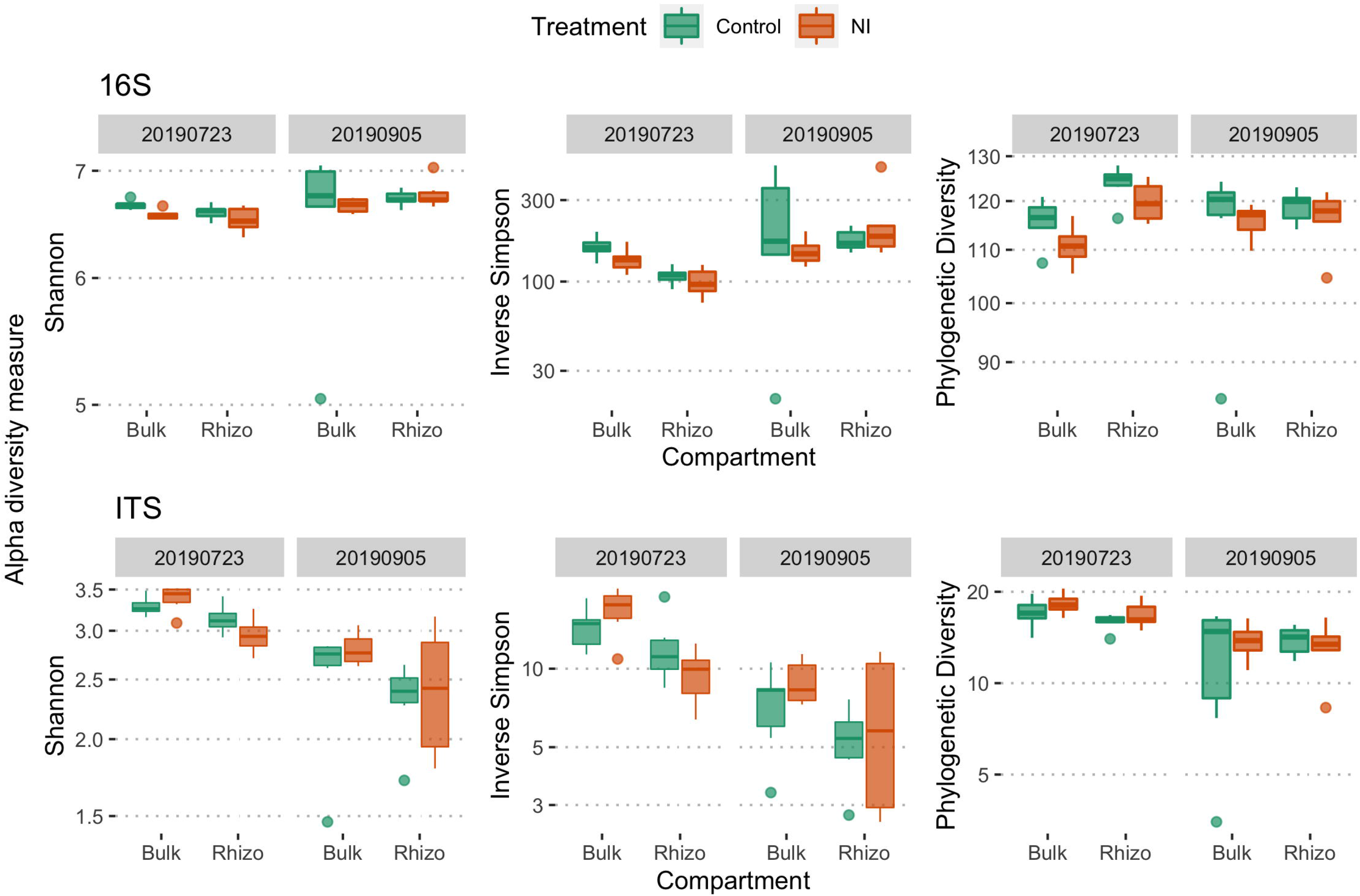
Alpha diversity of bacterial, archaeal (16S) and fungal (ITS) communities. Diversity indices were calculated using Shannon diversity index, Inverse Simpson index and Faith’s Phylogenetic Diversity (PD) across NI treatments and sampling dates. The boxplot shows quartile values per compartment (bulk soil and rhizosphere) and are colored by treatment and facetted by sampling date (2019-07-23, 2019-09-05).

At the phylum level, bacterial and archaeal communities were dominated, in descending order, by the phyla (mean relative abundance >1% across all samples) Thaumarchaeota, Verrucomicrobia, Proteobacteria, Actinobacteria, Nitrospirae, Acidobacteria, Firmicutes, Bacteroidetes and Gemmatimonadetes (Figure 2A), whereas fungal communities were dominated, in descending order, by the phyla (mean relative abundance >1% across all samples) Mortierellomycota, Ascomycota, Basidiomycota and Blastocladiomycota (Figure 2B).

**Figure 2:**
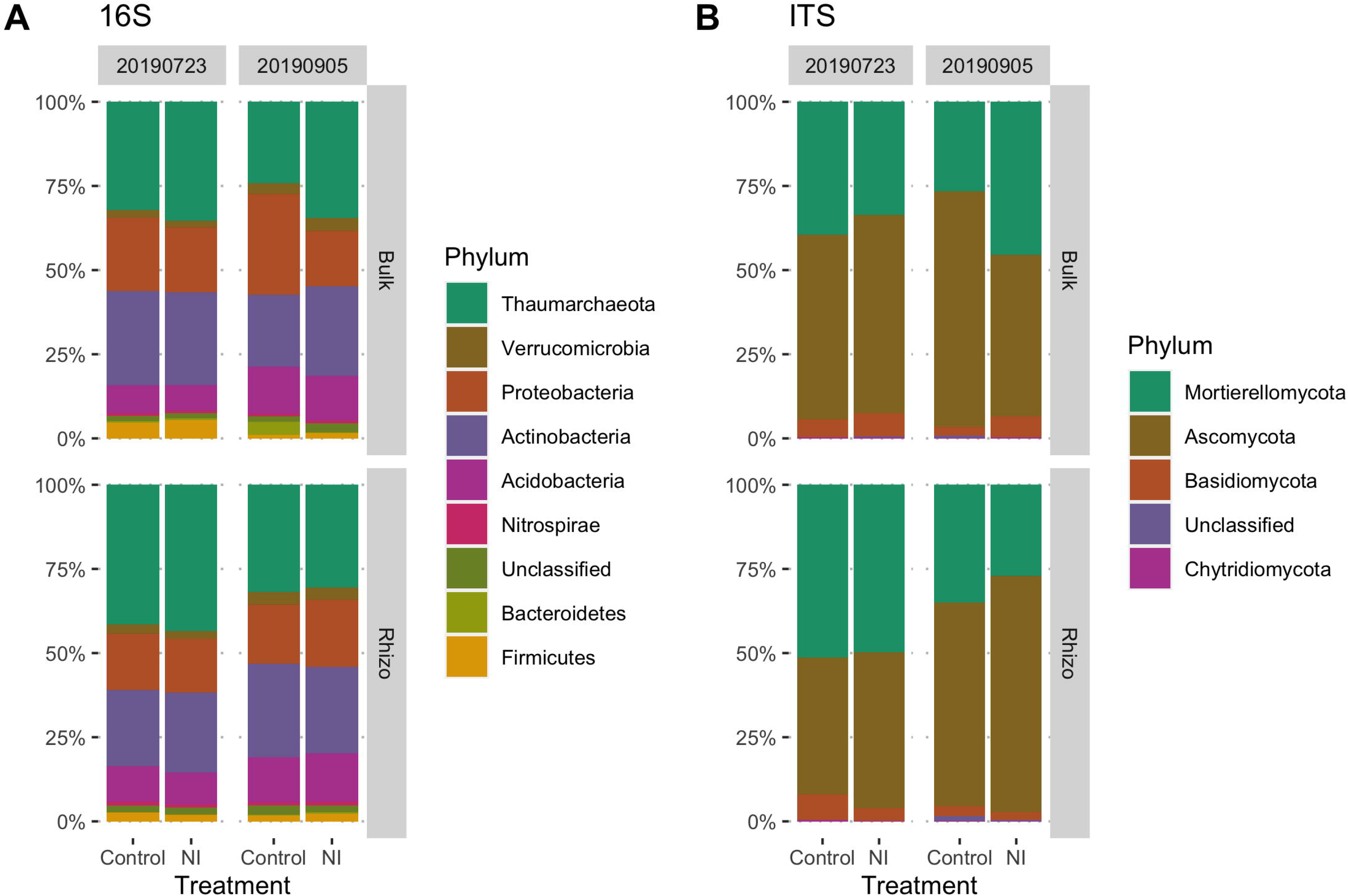
Bacterial (16S, A) and fungal (ITS, B) community composition on phylum level. The mean relative abundance above 1% was calculated based on Illumina amplicon sequencing of the 16S rRNA gene and the ITS region for bacteria and archaea, and fungi, respectively. Treatment: NI: Fertilizer with nitrapyrin, Control: Fertilizer without nitrapyrin.

Out of the nine bacterial and archaeal phyla and four fungal phyla listed above, nitrapyrin as a main effect only had a significant effect on the relative abundance of the archaeal phylum Thaumarchaeota with an increased relative abundance in the NI treatment (Table 2, Figure 3). In contrast, compartment and sampling date significantly affected the relative abundance a most bacterial, archaeal and fungal phyla (Table 2 and Figure 3). The interaction effect between the NI treatment and compartment was significant for Thaumarchaeota, Proteobacteria and Basidiomycota, whereas the interaction effect between nitrapyrin and sampling date was only significant for Nitrospirae. The interaction effect between compartment and sampling date was significant for Thaumarchaeota, Actinobacteria, Firmicutes, Mortierellomycota and Ascomycota. The three-way interaction effect was not significant for any of the major phyla tested.

**Table 2:** Three-way repeated measures ANOVA of the relative abundance of dominant phyla of bacterial, archaeal (16S) and fungal (ITS) communities in descending order of abundance.

| 16S Phyla | T |  | C |  | D |  | T x C |  | T x D |  | C x D |  | T x C x D |  |
| --- | --- | --- | --- | --- | --- | --- | --- | --- | --- | --- | --- | --- | --- | --- |
|  | F | P | F | P | F | P | F | P | F | P | F | P | F | P |
| Thaumarchaeota | 4.463 | **0.041** | 11.899 | **0.001** | 23.796 | **< 0.001** | 5.845 | **0.02** | 0.282 | 0.598 | 5.301 | **0.027** | 3.873 | 0.056 |
| Verrucomicrobia | 0.216 | 0.645 | 3.353 | 0.075 | 42.408 | **< 0.001** | 1.456 | 0.235 | 2.848 | 0.099 | 0.341 | 0.563 | 1.353 | 0.252 |
| Proteobacteria | 2.233 | 0.143 | 6.288 | **0.016** | 2.218 | 0.144 | 4.912 | **0.032** | 0.917 | 0.344 | 0.047 | 0.829 | 3.643 | 0.064 |
| Actinobacteria | 0.571 | 0.454 | 1.162 | 0.288 | 0.019 | 0.89 | 0.581 | 0.45 | 0.292 | 0.592 | 17.476 | **< 0.001** | 2.934 | 0.094 |
| Nitrospirae | 0.041 | 0.841 | 17.094 | **< 0.001** | 0.067 | 0.797 | 0.353 | 0.556 | 4.91 | **0.032** | 1.626 | 0.21 | 0.751 | 0.391 |
| Acidobacteria | 0.314 | 0.578 | 4.703 | **0.036** | 50.902 | **< 0.001** | 0.095 | 0.759 | 0.838 | 0.366 | 1.952 | 0.17 | 0.001 | 0.979 |
| Firmicutes | 0.828 | 0.368 | 10.89 | **0.002** | 75.806 | **< 0.001** | 2.044 | 0.161 | 1.378 | 0.247 | 19.477 | **< 0.001** | 1.766 | 0.191 |
| Bacteroidetes | 1.346 | 0.253 | 2.812 | 0.101 | 0.548 | 0.463 | 2.205 | 0.145 | 1.075 | 0.306 | 0.127 | 0.723 | 1.889 | 0.177 |
| Gemmatimonadetes | 1.605 | 0.213 | 0.897 | 0.349 | 10.954 | **0.002** | 0.323 | 0.573 | 0.386 | 0.538 | 1.292 | 0.262 | 0.817 | 0.371 |
| **ITS Phyla** |  |  |  |  |  |  |  |  |  |  |  |  |  |  |
| Mortierellomycota | 0.037 | 0.849 | 1.505 | 0.227 | 12.566 | **0.001** | 1.561 | 0.219 | 2.924 | 0.095 | 7.27 | **0.01** | 2.407 | 0.129 |
| Ascomycota | 0.081 | 0.777 | 0.897 | 0.349 | 17.854 | **< 0.001** | 2.676 | 0.11 | 4.004 | 0.052 | 7.177 | **0.011** | 2.161 | 0.149 |
| Basidiomycota | 0.044 | 0.836 | 2.415 | 0.128 | 25.571 | **< 0.001** | 6.092 | **0.018** | 3.15 | 0.084 | 0.076 | 0.784 | 0.675 | 0.416 |
| Blastocladiomycota | 1.623 | 0.21 | 0.531 | 0.47 | 0.564 | 0.457 | 0.378 | 0.542 | 0.748 | 0.392 | 0.995 | 0.325 | 1.235 | 0.273 |

T: Treatment with nitrapyrin and control without nitrapyrin. C: Compartment (Bulk soil and Rhizosphere) D: sampling dates (2019-07-23, 2019-09-05). *P*-values marked in bold indicate statistically significant difference with $P < 0.05$ .

**Figure 3:**
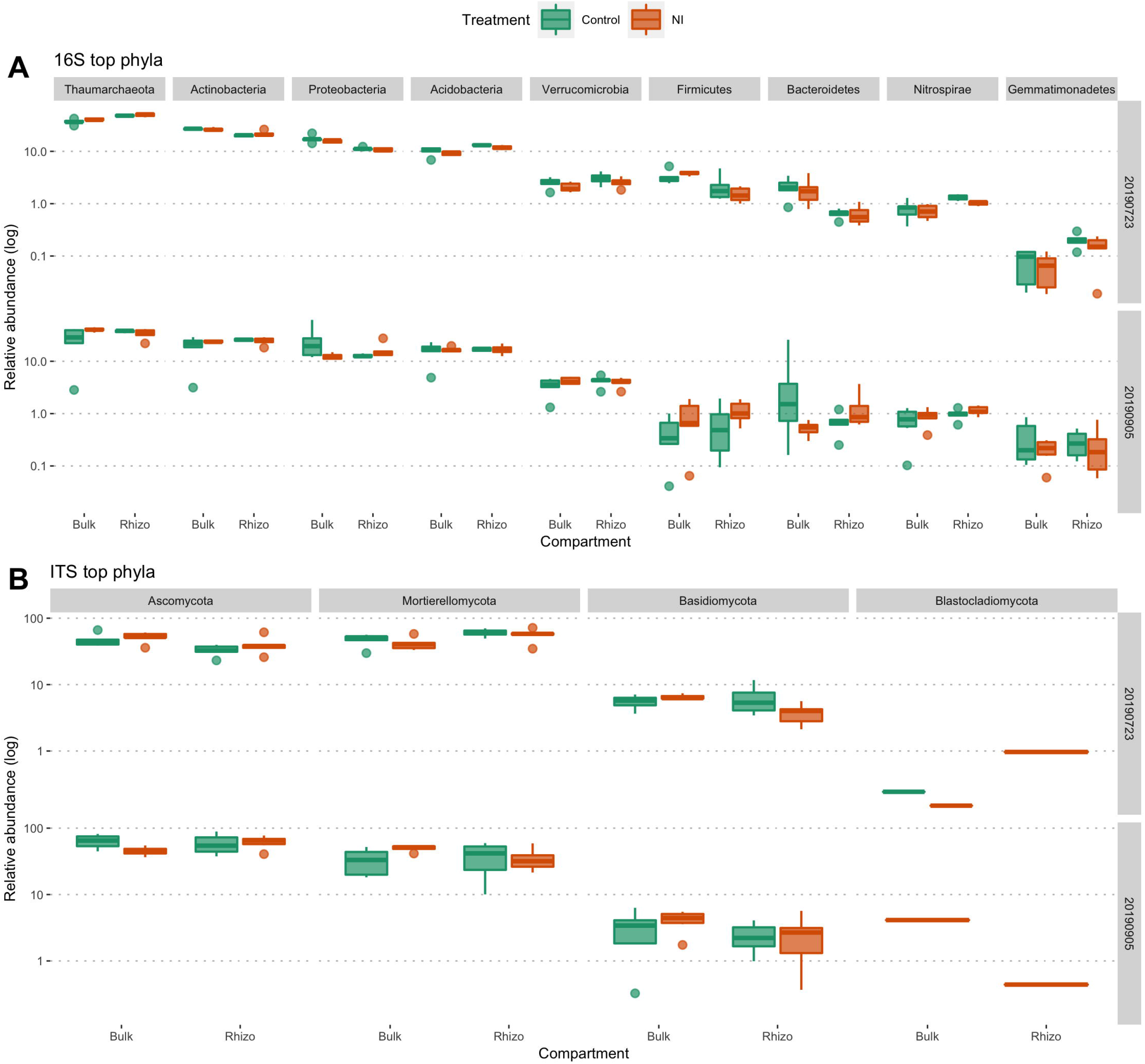
Variation of relative abundance of dominant bacterial, archaeal (16S) and fungal (ITS) phyla across treatments. Treatment: NI: Fertilizer with nitrapyrin, Control: Fertilizer without nitrapyrin, compartment (bulk soil and rhizosphere) and sampling dates (2019-07-23, 2019-09-05).

At the genus level, we focused our analyses on the 10 relatively most abundant genera for the archaea and bacteria, and for the fungi (listed in Table 3). The main effect of NI treatment was only significant for the genus *Nitrososphaera* (Table 3, Figure 4). In contrast, the main effect of compartment was significant for 5 out of 10 bacterial and archaeal genera and for one fungal genus (Table 3), whereas the main effect of sampling date was significant for 6 out of 10 bacterial and archeal genera and 7 out of 10 fungal genera (Table 3). The interaction effect between nitrapyrin addition and compartment was significant for *Nitrososphaera*, *Gaiella*, *Rhodoplanes*, *Gliomastix* and *Ganoderma*, whereas the interaction effect between nitrapyrin and sampling date was not significant for any of the most abundant genera (Table 3). The interaction effect between compartment and sampling date was significant for *Nitrososphaera*, *Solirubrobacter*, *Hyphomicrobium, Rhodoplanes*, *Mortierella* and *Thelonectria* (Table 3). Furthermore, the three-way interaction was significant for *Gaiella*, *Solirubrobacter*, *Hyphomicrobium* and *Gliomastix* (Table 3).

**Table 3:**
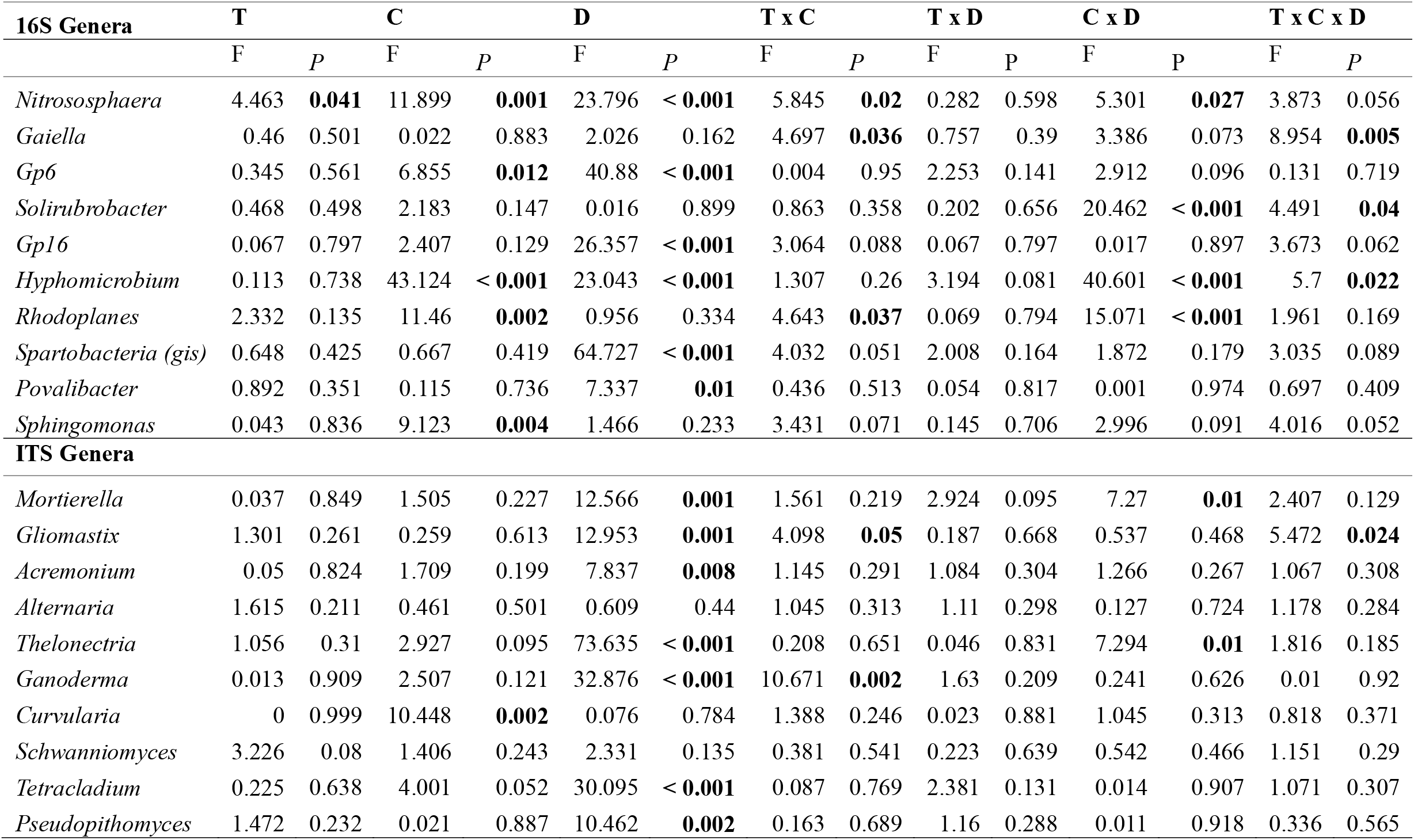
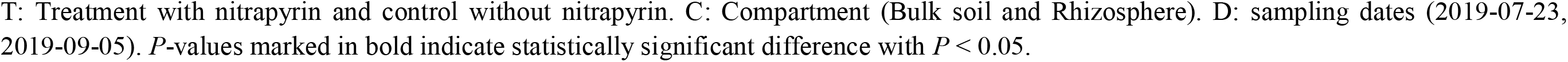
Three-way repeated measures ANOVA of the relative abundance of dominant genera of bacterial, archaeal (16S) and fungal (ITS) communities in descending order of abundance.

| 16S Genera | T |  | C |  | D |  | T x C |  | T x D |  | C x D |  | T x C x D |  |
| --- | --- | --- | --- | --- | --- | --- | --- | --- | --- | --- | --- | --- | --- | --- |
|  | F | P | F | P | F | P | F | P | F | P | F | P | F | P |
| *Nitrososphaera* | 4.463 | **0.041** | 11.899 | **0.001** | 23.796 | **< 0.001** | 5.845 | **0.02** | 0.282 | 0.598 | 5.301 | **0.027** | 3.873 | 0.056 |
| *Gaiella* | 0.46 | 0.501 | 0.022 | 0.883 | 2.026 | 0.162 | 4.697 | **0.036** | 0.757 | 0.39 | 3.386 | 0.073 | 8.954 | **0.005** |
| *Gp6* | 0.345 | 0.561 | 6.855 | **0.012** | 40.88 | **< 0.001** | 0.004 | 0.95 | 2.253 | 0.141 | 2.912 | 0.096 | 0.131 | 0.719 |
| *Solirubrobacter* | 0.468 | 0.498 | 2.183 | 0.147 | 0.016 | 0.899 | 0.863 | 0.358 | 0.202 | 0.656 | 20.462 | **< 0.001** | 4.491 | **0.04** |
| *Gp16* | 0.067 | 0.797 | 2.407 | 0.129 | 26.357 | **< 0.001** | 3.064 | 0.088 | 0.067 | 0.797 | 0.017 | 0.897 | 3.673 | 0.062 |
| *Hyphomicrobium* | 0.113 | 0.738 | 43.124 | **< 0.001** | 23.043 | **< 0.001** | 1.307 | 0.26 | 3.194 | 0.081 | 40.601 | **< 0.001** | 5.7 | **0.022** |
| *Rhodoplanes* | 2.332 | 0.135 | 11.46 | **0.002** | 0.956 | 0.334 | 4.643 | **0.037** | 0.069 | 0.794 | 15.071 | **< 0.001** | 1.961 | 0.169 |
| *Spartobacteria (gis)* | 0.648 | 0.425 | 0.667 | 0.419 | 64.727 | **< 0.001** | 4.032 | 0.051 | 2.008 | 0.164 | 1.872 | 0.179 | 3.035 | 0.089 |
| *Povalibacter* | 0.892 | 0.351 | 0.115 | 0.736 | 7.337 | **0.01** | 0.436 | 0.513 | 0.054 | 0.817 | 0.001 | 0.974 | 0.697 | 0.409 |
| *Sphingomonas* | 0.043 | 0.836 | 9.123 | **0.004** | 1.466 | 0.233 | 3.431 | 0.071 | 0.145 | 0.706 | 2.996 | 0.091 | 4.016 | 0.052 |
| **ITS Genera** |  |  |  |  |  |  |  |  |  |  |  |  |  |  |
| *Mortierella* | 0.037 | 0.849 | 1.505 | 0.227 | 12.566 | **0.001** | 1.561 | 0.219 | 2.924 | 0.095 | 7.27 | **0.01** | 2.407 | 0.129 |
| *Gliomastix* | 1.301 | 0.261 | 0.259 | 0.613 | 12.953 | **0.001** | 4.098 | **0.05** | 0.187 | 0.668 | 0.537 | 0.468 | 5.472 | **0.024** |
| *Acremonium* | 0.05 | 0.824 | 1.709 | 0.199 | 7.837 | **0.008** | 1.145 | 0.291 | 1.084 | 0.304 | 1.266 | 0.267 | 1.067 | 0.308 |
| *Alternaria* | 1.615 | 0.211 | 0.461 | 0.501 | 0.609 | 0.44 | 1.045 | 0.313 | 1.11 | 0.298 | 0.127 | 0.724 | 1.178 | 0.284 |
| *Thelonectria* | 1.056 | 0.31 | 2.927 | 0.095 | 73.635 | **< 0.001** | 0.208 | 0.651 | 0.046 | 0.831 | 7.294 | **0.01** | 1.816 | 0.185 |
| *Ganoderma* | 0.013 | 0.909 | 2.507 | 0.121 | 32.876 | **< 0.001** | 10.671 | **0.002** | 1.63 | 0.209 | 0.241 | 0.626 | 0.01 | 0.92 |
| *Curvularia* | 0 | 0.999 | 10.448 | **0.002** | 0.076 | 0.784 | 1.388 | 0.246 | 0.023 | 0.881 | 1.045 | 0.313 | 0.818 | 0.371 |
| *Schwanniomyces* | 3.226 | 0.08 | 1.406 | 0.243 | 2.331 | 0.135 | 0.381 | 0.541 | 0.223 | 0.639 | 0.542 | 0.466 | 1.151 | 0.29 |
| *Tetracladium* | 0.225 | 0.638 | 4.001 | 0.052 | 30.095 | **< 0.001** | 0.087 | 0.769 | 2.381 | 0.131 | 0.014 | 0.907 | 1.071 | 0.307 |
| *Pseudopithomyces* | 1.472 | 0.232 | 0.021 | 0.887 | 10.462 | **0.002** | 0.163 | 0.689 | 1.16 | 0.288 | 0.011 | 0.918 | 0.336 | 0.565 |

T: Treatment with nitrapyrin and control without nitrapyrin. C: Compartment (Bulk soil and Rhizosphere). D: sampling dates (2019-07-23, 2019-09-05). *P*-values marked in bold indicate statistically significant difference with $P < 0.05$ .

**Figure 4:**
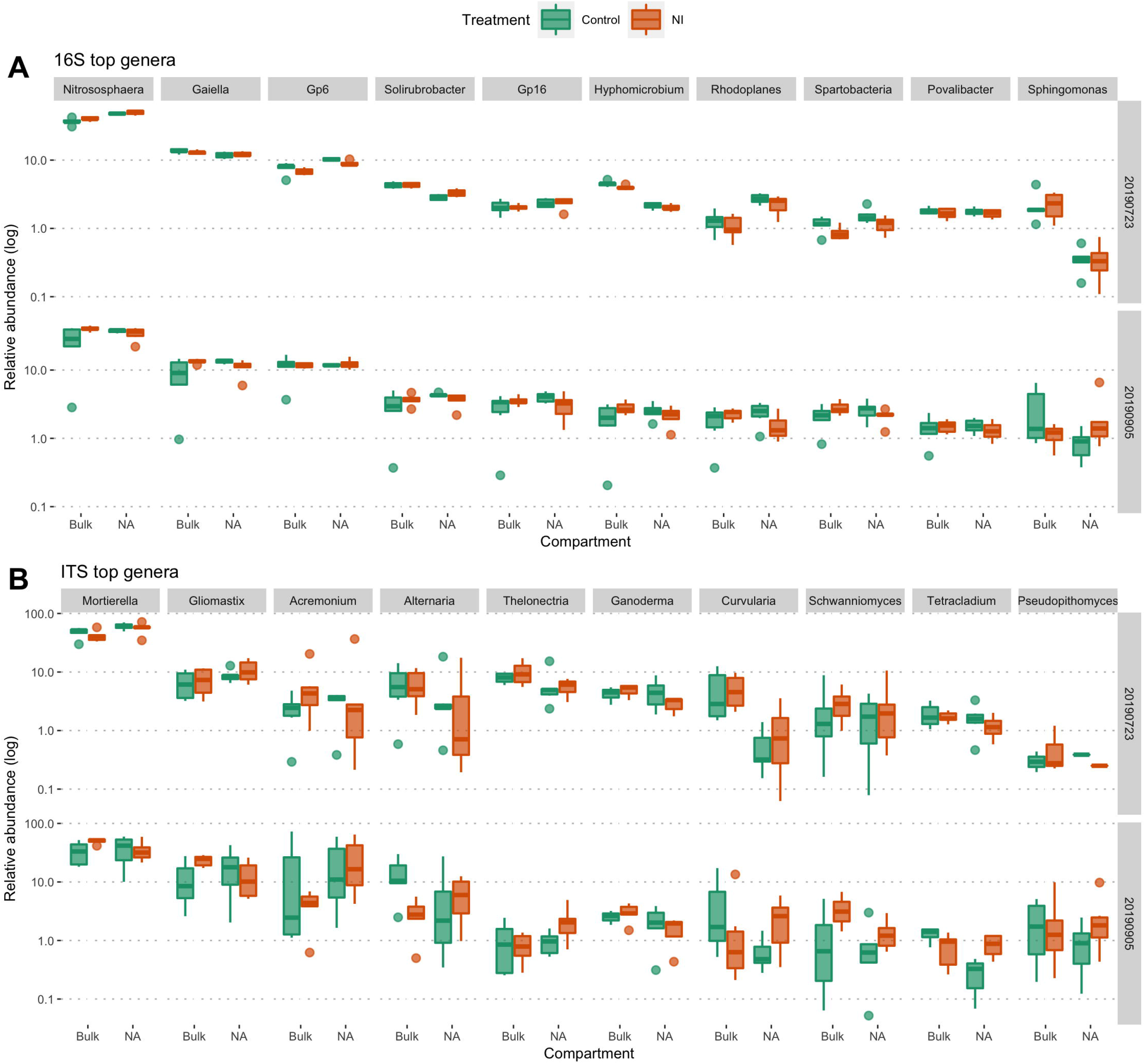
Variation of relative abundance of dominant bacterial, archaeal (16S) and fungal (ITS) genera across treatments. Treatment: NI: Fertilizer with nitrapyrin, Control: Fertilizer without nitrapyrin, compartment (bulk soil and rhizosphere) and sampling dates (2019-07-23, 2019-09-05).

The overall structures of bacterial and archaeal, and fungal communities across sampling dates are shown in the principal coordinates analysis (PCoA) ordinations in Figure 5. For both, 16S and ITS ordinations, the first axis clearly separated between the rhizosphere and bulk soil samples only for the first sampling date (Figure 5). No clearly visible effect of nitrapyrin could be observed in the ordination plots. The Permanova analyses confirmed this visual interpretation for both archaeal, bacterial and fungal communities, with significant main effects for compartment and sampling date, but not for nitrapyrin addition (Table 4). Additionally, for both communities, the interaction effects between nitrapyrin and compartment and between sampling date and compartment were significant, whereas the three-way interaction effect was also significant for bacteria and archaea (Table 4).

**Table 4:** PERMANOVA of bacterial, archaeal (16S) and fungal (ITS) community structure for the effects of Treatment (treatment with nitrapyrin and control without nitrapyrin), Compartment (Bulk soil and Rhizosphere), Date (2019-07-23, 2019-09-05) and interactions. The significance of observed R was assessed by permuting (9999 permutations). *P*-values marked in bold indicate statistically significant difference with *P* < 0.05.

|  | **16S** |  | **ITS** |  |
| --- | --- | --- | --- | --- |
| | $R^2$ | *P* | $R^2$ | *P* |
| Treatment | 0.019 | 0.244 | 0.016 | 0.422 |
| Compartment | 0.05 | **0.001** | 0.035 | **0.020** |
| Date | 0.124 | **0.001** | 0.187 | **0.001** |
| Treatment × Compartment | 0.041 | **0.004** | 0.031 | **0.048** |
| Treatment × Date | 0.015 | 0.464 | 0.016 | 0.408 |
| Date × Compartment | 0.057 | **0.002** | 0.033 | **0.038** |
| Treatment × Date × Compartment | 0.036 | **0.019** | 0.029 | 0.076 |

Treatment: treatment with nitrapyrin and control without nitrapyrin. C: Compartment (Bulk soil and Rhizosphere). Date: sampling dates (2019-08-20, 2019-09-05). *P*-values marked in bold indicate statistically significant difference with $P < 0.05$ .

**Figure 5:**
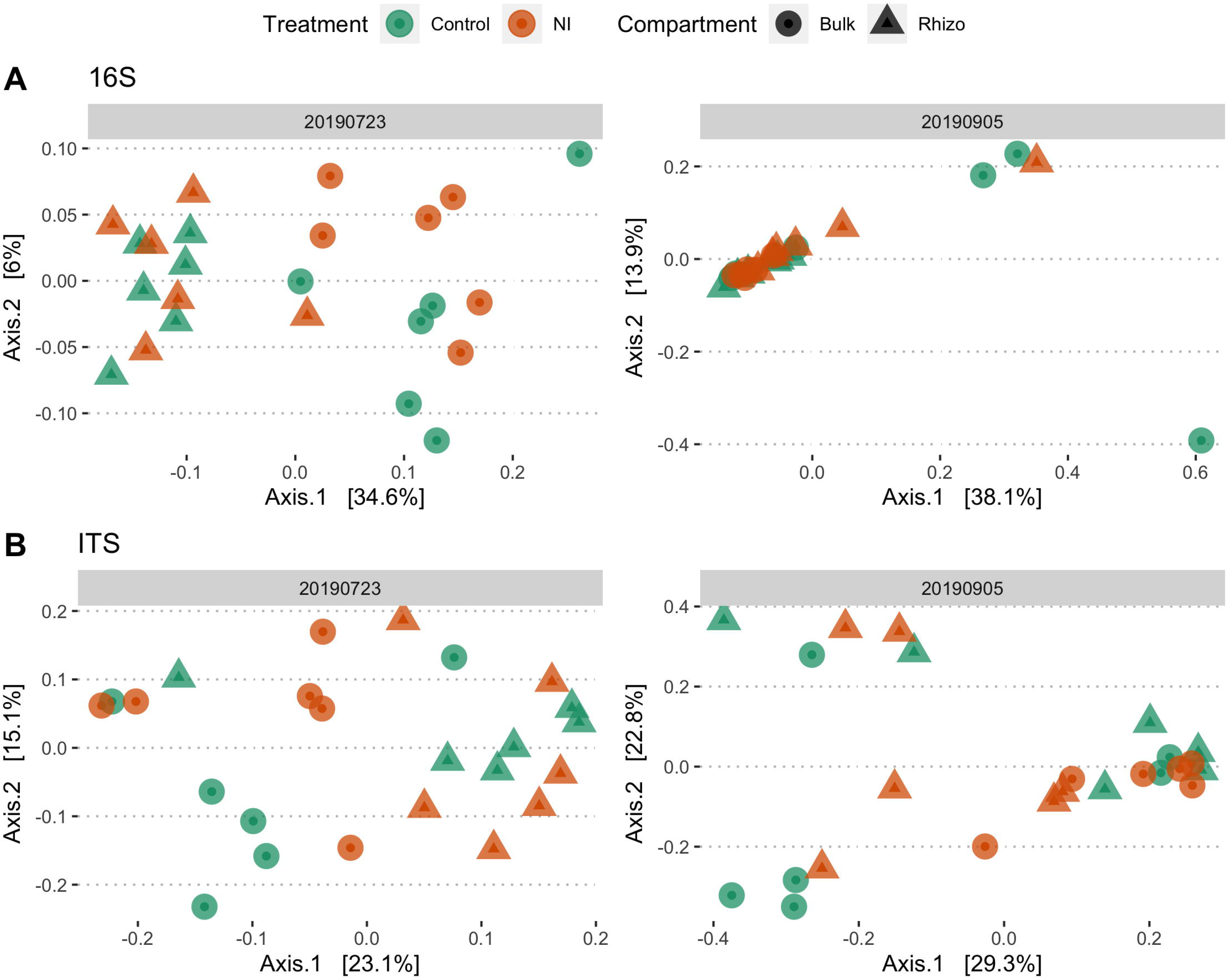
Principal Coordinates Analysis (PCoA) of the bacterial, archaeal (16S) and fungal (ITS) community composition based on Bray-Curtis dissimilarity. The first and second axes of the 16S ordination explained 34.6% and 6% of the variance for sampling date 2019-07-23, and 38.1% and 13.9% of the variance for sampling date 2019-09-05. For ITS, the first and second axes of explained 23.1% and 15.1% of the variance for sampling date 2019-07-23, and 29.3% and 22.8% of the variance for sampling date 2019-09-05. Treatment: NI: Fertilizer with nitrapyrin, Control: Fertilizer without nitrapyrin, compartment (bulk soil and rhizosphere).

### Abundance of functional genes of the N cycle

The abundances of the AOA-*amoA*, AOB-*amoA*, *nirK*, *nifH* and *nirS* genes as well as AOA to AOB ratio were not individually affected by the main effect of the NI treatment, but were all strongly affected by the main effect of sampling date (Table 5). However, the interaction term between the NI treatment and sampling date, between the compartment and sampling date, as well as the three-way interaction effect were highly significant for all the genes and for the AOA to AOB ratio measured (Table 5). Specifically, the abundance of *amoA*-AOA, *nirK* and *nifH* significantly increased over time across all treatments, whereas the abundance of *nirS* increased over time only in NI treatments in rhizosphere soil (Figure 6). In contrast, the abundance of *amoA*-AOB significantly decreased over time across all treatments (Figure 6). AOA to AOB ratios increased over time in both compartments, with a higher ratio in bulk soil as compared to the rhizosphere (Figure 6).

**Table 5:** Kruskal-Wallis test of the abundance of functional genes involved in the nitrogen cycle.

|  | **T** |  | **C** |  | **D** |  | **T x C** |  | **T x D** |  | **C x D** |  | **T x C x D** |  |
| --- | --- | --- | --- | --- | --- | --- | --- | --- | --- | --- | --- | --- | --- | --- |
| | $\chi^2$ | *P* | $\chi^2$ | *P* | $\chi^2$ | *P* | $\chi^2$ | *P* | $\chi^2$ | *P* | $\chi^2$ | *P* | $\chi^2$ | *P* |
| *amoA*-AOA | 1.021 | 0.312 | 1.150 | 0.284 | 21.716 | < **0.001** | 2.205 | 0.531 | 22.741 | < **0.001** | 25.724 | < **0.001** | 27.049 | < **0.001** |
| *amoA*-AOB | 0.435 | 0.509 | 0.17 | 0.68 | 22.491 | < **0.001** | 2.24 | 0.524 | 22.954 | < **0.001** | 28.582 | < **0.001** | 30.816 | < **0.001** |
| AOA:AOB ratio | 0.188 | 0.665 | 0.043 | 0.837 | 27.215 | < **0.001** | 1.71 | 0.635 | 27.445 | < **0.001** | 32.125 | < **0.001** | 28.16 | < **0.001** |
| *nirK* | 0.002 | 0.962 | 1.728 | 0.189 | 32 | < **0.001** | 1.774 | 0.621 | 32.683 | < **0.001** | 34.569 | < **0.001** | 35.624 | < **0.001** |
| *nirS* | 0.051 | 0.821 | 2.521 | 0.112 | 14.551 | < **0.001** | 4.314 | 0.23 | 19.38 | < **0.001** | 21.324 | < **0.001** | 28.16 | < **0.001** |
| *nifH* | 1.734 | 0.188 | 0.322 | 0.57 | 32.87 | < **0.001** | 2.121 | 0.548 | 36.287 | < **0.001** | 33.454 | < **0.001** | 37.265 | < **0.001** |

T: Treatment with nitrapyrin and control without nitrapyrin. C: Compartment (Bulk soil and Rhizosphere). Date: sampling dates (2019-08-20, 2019-09-05). *P*-values marked in bold indicate statistically significant difference with $P < 0.05$ .

**Figure 6:**
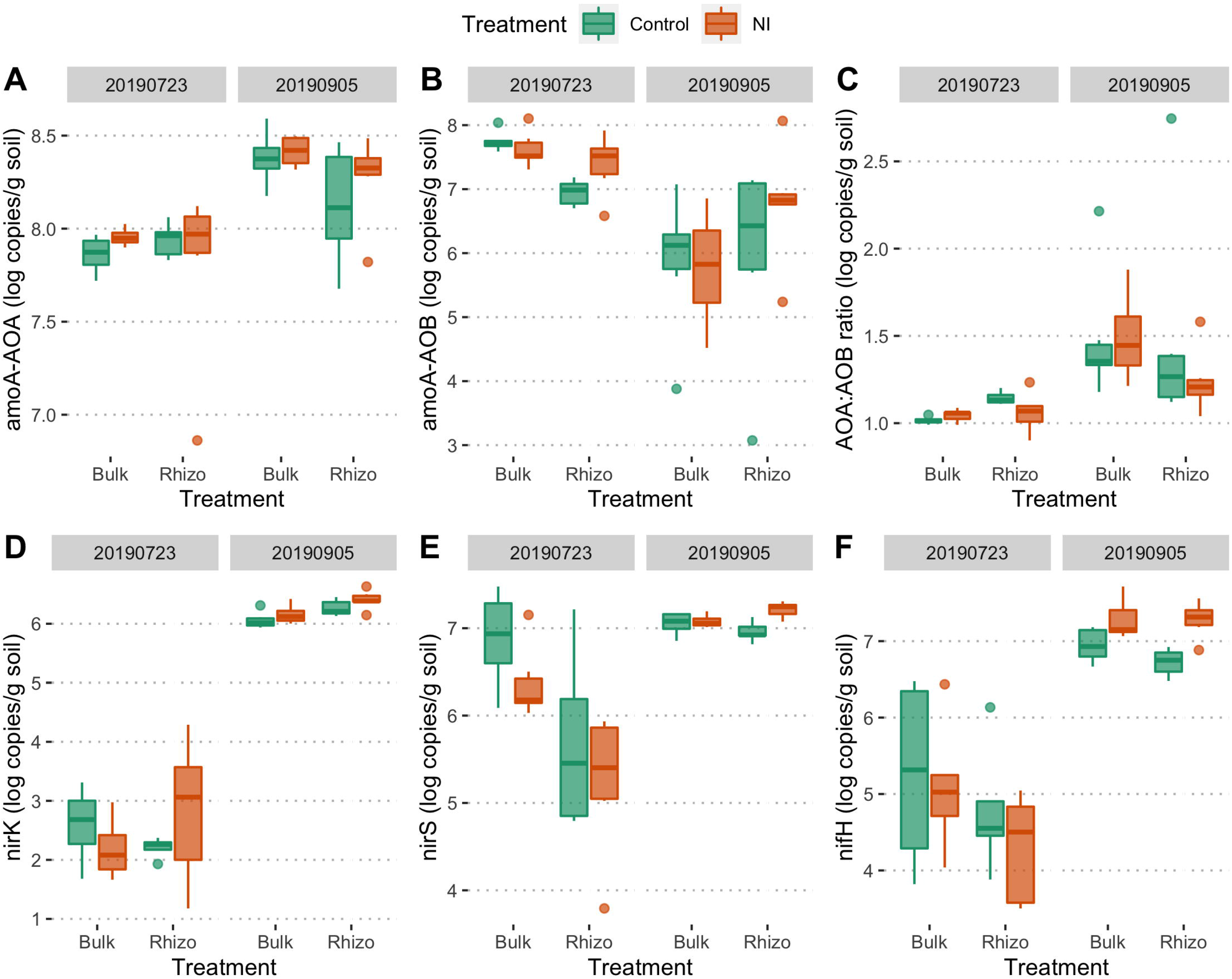
Abundance of the archaeal (A) and bacterial (B) ammonia monooxygenase A subunit (*amoA*-AOA and *amoA*-AOB, respectively) genes, AOA:AOB ratio (C), *nirK* (D), *nirS* (E) and *nifH* (F) genes. Abundances (log copies/g soil) were determined by quantitative PCR (qPCR) across treatments, compartment and sampling dates. The boxplot shows quartile values colored by treatments. Treatment: NI: Fertilizer with nitrapyrin, Control: Fertilizer without nitrapyrin, compartment (bulk soil and rhizosphere) and sampling dates (2019-07-23, 2019-09-05).

### Volatile organic compound (VOC) composition

The overall composition of rhizosphere VOCs across treatment and sampling date can be visualized through the first two components of the principal component analysis (PCA) (Figure 7). In Permanova tests, VOC profiles were not significantly affected by the NI treatment but differed significantly among sampling date (Table 6, Figure 7). No individual compounds were found to be significantly different between the control and the NI treatment. However, thirteen compounds showing significant (P<0.001) differences between sampling dates were identified (Figure 8, Table S3). These compounds, ordered by decreasing significance values in Table S3, were tentatively annotated as members of the chemical classes of terpene, ester, alcohol and alkene. The monoterpene alpha-pinene showed the most significant respond to sampling date, with a large increase over time in both the NI and control treatments (Figure 8B).

**Table 6:** PERMANOVA of rhizosphere VOC composition for the effects of treatment, date and interactions. The significance of observed R was assessed by permuting (9999 permutations).

| | $R^2$ | $P$ |
| --- | --- | --- |
| Treatment | 0.013 | 0.384 |
| Date | 0.665 | **0.001** |
| Treatment × Date | 0.016 | 0.304 |

Treatment: treatment nitrapyrin and control without nitrapyrin. Date: sampling dates (2019-08-20, 2019-09-05). $P$ -values marked in bold indicate statistically significant difference with $P < 0.05$ .

**Figure 7:**
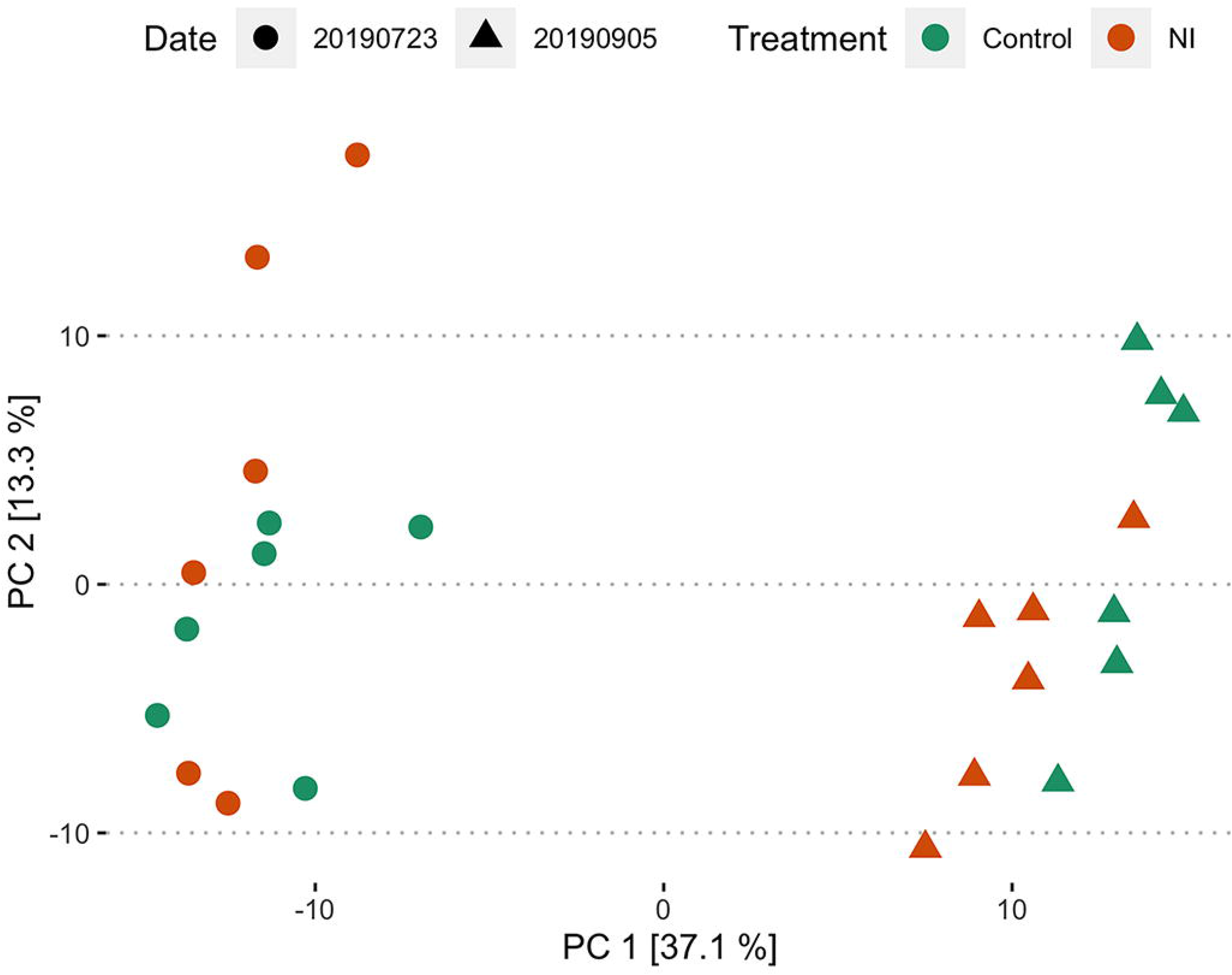
(A) Principal Component Analysis (PCA) of the rhizosphere VOC composition. The first and second axes of the ordination explained 37.1% and 13.3% of the variance. Treatment: NI: Fertilizer with nitrapyrin, Control: Fertilizer without nitrapyrin, and sampling dates (2019-07-23, 2019-09-05).

**Figure 8:**
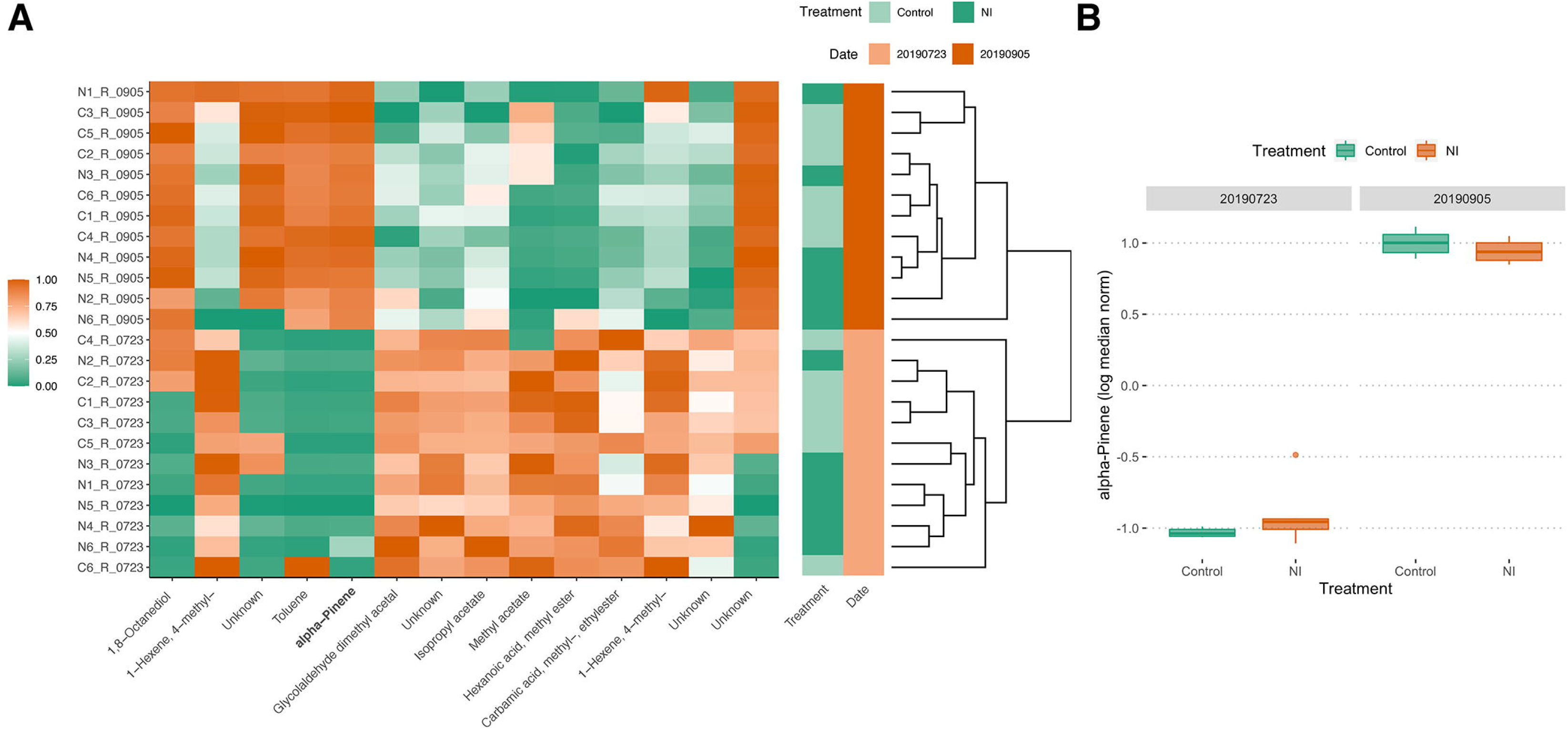
Heatmap of VOCs emitted based Euclidian distance-clustering of samples based on sampling date. **(A).** ‘Unknown’ indicates no match was found in the NIST library 2014 V2.20, or based on mass spectra, retention time and retention index (Table S3). Treatment: NI: Fertilizer with nitrapyrin, Control: Fertilizer without nitrapyrin, and sampling dates (2019-07-23, 2019-09-05). **Alpha-pinene concentration across treatment and sampling dates (B)**. Treatment: NI: Fertilizer with nitrapyrin, Control: Fertilizer without nitrapyrin, and sampling dates (2019-07-23, 2019-09-05). Values are depicted as log median normalised values.

## Discussion

The inhibitory effect of nitrapyrin on soil microbial communities are not entirely understood. It remains unresolved whether this nitrification inhibitor (NI) leads to direct or indirect shifts in the overall soil microbial community structure and functioning. The effects of adding various NIs together with N-containing fertilizers have been shown to range from no changes in the community structure of ammonia-oxidizers and in the abundances of total bacteria (24, 65) to only minimal and transient changes on the overall microbial community structure, depending on the compartment (66). In contrast to these findings, we found here widespread effects of nitrapyrin on soil and wheat rhizosphere microbial communities across different sampling dates. For instance, many of the major phyla were affected by the nitrapyrin addition, though this effect was often variable at the different sampling dates, or in the different soil compartments. It is difficult to ascertain if these shifts were caused directly by nitrapyrin (e.g. off-target toxic effects) or indirectly through the reduced activities of ammonia-oxidizers. Nevertheless, our results suggest, in agreement with our hypothesis, that the effects of nitrification inhibitors might have far reaching but difficult to predict effects on soil functioning.

Nitrapyrin also had significant effects on its expected targets, the ammonia-oxidizers. Nitrapyrin is generally thought to be a strong inhibitor of AOBs, with limited efficiency for AOA (67, 68). We were thus expecting a decrease in AOB for the nitrapyrin plots, and potentially an increase in AOA because of the lowered competition from AOB. For AOA, we did find an increase in both the qPCR and 16S rRNA gene data. The dominant genera found in our samples, the AOA *Nitrososphaera*, and the associated phylum (Thaumarchaeota) significantly increased in relative abundance following the addition of nitrapyrin. This shift in the relative abundance based on 16S rRNA sequencing was confirmed by the real-time PCR quantification of the archaeal *amoA* gene. In this case, the increase was dependent on the sampling date, and was more important at the first sampling date. Surprisingly, similar trends were observed for the abundance of AOB *amoA* gene in the rhizosphere of wheat, suggesting a weak efficiency of the nitrapyrin on its target group. Previous studies on the effect of nitrapyrin on the AOA to AOB ratio and its effect on nitrification rates have reported inconclusive results, with nitrapyrin affecting either AOA, AOB or both (21–24).

In the rhizosphere of wheat, the increase following nitrapyrin addition was stronger for the AOB, but AOA still predominated in most samples at both sampling date. This is in line with previous work that reported that AOA and not AOB control nitrification in soil (7, 69), although some studies reported otherwise (70). AOA are also thought to be favoured over AOB in the rhizosphere due to a better competitive ability against plants for ammonia (71). Interestingly, the AOA to AOB ratio was not stable through time as AOA *amoA* increased with time and AOB *amoA* decreased with time. This suggests that, next to variations between fields with different soil physico-chemical characteristics, the relative dominance of AOA and AOB might vary through the seasons within a field.

For most results, the effect of nitrapyrin depended on sampling date or compartment. Sampling date was the overarching factor affecting the abundance of all N-cycle genes. The effect of nitrapyrin was often clearer and stronger at the first sampling date. This is not surprising as the nitrapyrin was applied twice, in June and July, and the first sampling date was two weeks after the second application, whereas the last sampling date was in September. Nitrapyrin has a limited residence time in soil (72), and it was probably not present in sufficient quantities to cause any effect other than legacy effects at the last sampling date.

Apart from the specific effect on ammonia-oxidizers, the nitrapyrin treatment, among and overarching effect of compartment and sampling date, affected the relative abundance of many bacterial, archaeal and fungal taxa. For example, Proteobacteria were affected by the nitrapyrin treatment depending on the compartment, with a decrease in relative abundance in the bulk soil as compared to the rhizosphere in the nitrapyrin treatment. Moreover, the nitrite-oxidizing bacterial phylum Nitrospirae that often occurs in close association with AOB or AOA (73) was affected by the nitrapyrin treatment depending on the sampling date, with an increase in relative abundance in the nitrapyrin treatment over time. The only fungal phylum that was affected by the nitrapyrin treatment dependent on the compartment was Basidiomycota, which contain many denitrifiers that play key roles in the N cycle (74). Similarly, genes related to denitrification were affected by nitrapyrin in interaction with sampling date, most probably through changes in the availability of nitrate caused by alterations in nitrification rates. These results confirm our hypothesis that nitrapyrin alters the relative abundance of non-target bacteria, archaea, fungi and associated nitrogen-cycle functions.

In order to measure functional changes beyond the abundance of functional genes, we measured detectable shifts in the metabolomics activities of the rhizosphere microbes and plant metabolites, namely VOCs. In contrast, nitrapyrin did not significantly influence the VOC metabolic profile in the rhizosphere of wheat. This suggests that the changes observed in the microbial community composition, structure and functions at the DNA level did not result in major shifts in the metabolic activities of the rhizosphere microbes and plant roots. Although we cannot exclude that other classes of low-molecular-weight metabolites, especially trace gases involved in the N-cycle, might have been affected by nitrapyrin, our results are suggesting that nitrapyrin might not cause large changes in the general microbial activities in the rhizosphere of wheat. This could be related to the high functional redundancy typically found in agricultural soils that is responsible for maintaining ecosystem functioning (75).

VOCs emitted by microbes in the rhizosphere are key metabolites driving microbial interactions and influencing biogeochemical processes in soil, including the N-cycle (25, 76, 77). For example, bacterial VOCs have been shown to reduce the abundance of *nifH* and *nirS* genes (78), and inhibit nitrification (79). Although, in our study, we did not find a significant direct effect of the nitrapyrin treatment on VOC profiles, they clearly differed according to sampling date. This could be linked to a few VOCs that were different among sampling dates. Interestingly, alpha-pinene was identified as the most significantly different compound between the two dates, with a sharp increase at the second sampling date. Alpha-pinene, amongst other monoterpenes has been previously shown to inhibit nitrification by targeting the ammonia monooxygenase enzyme (80, 81), which could explain part of the large decrease in AOB *amoA* abundance at the second sampling date. These findings contrast a previous study that linked bacterial VOC emission to an increase in AOB *amoA* gene abundance, however could not identify terpenes responsible for the observed effect (78). Our results suggest that AOB might be more susceptible to the inhibitory effects of alpha-pinene than AOA. Generally, VOCs act in a synergistic and concentration-depend manner (82). Due to the use of a passive method for trapping VOCs that has limitations in detecting all VOCs present (83), potential other synergistic VOCs could not be identified. Moreover, the concentrations at which the inhibition in soil takes place remain unknown. What is known, however, is that the inhibition of nitrification by monoterpenes has substantial ecological implications (84). It has been already suggested that if the inhibitory activity of monoterpenes like alpha-pinene approaches that of nitrapyrin, then ecosystem-level processes like the N-cycle could be controlled by secondary plant products present in levels of a few parts per million (80). Generally, alpha-pinene alongside other monoterpenes is synthesized in foliage of many plant species, but can also be produced by microorganisms in soil where they regulate microbial activity and nutrient availability (85). As these VOCs were measured in the rhizosphere, it is plausible that the microbial community in the rhizosphere at least partly produced alpha-pinene. At this point, however, it remains unclear what ecological role alpha-pinene plays in the interaction between the plant, soil microbes and nitrifiers and what result it might have on the N-cycle.

In summary, we showed that, in agreement with our hypothesis, nitrapyrin addition resulted in far reaching changes in non-target microbial communities and in the abundance of N-cycle functional genes. However, these changes were not reflected in the metabolomic profiles of the rhizosphere, suggesting that nitrapyrin did not affect soil functionality. Additionally, nitrapyrin affected the abundance of AOA and AOB, but positively. Nitrapyrin effects were also constrained by sampling date and by the plant compartment sampled, suggesting an interaction with environmental conditions.

## Acknowledgements

This study was supported by a FRQNT Team Grant (2019-PR-254256). PG acknowledges NWO (VIDI 864.11.015) for financial support. We are grateful for the support of Liliana Quiza in the laboratory work on functional genes, and Hans Zweers and Maria Hundscheidt for preparing the PDMS tubes and measuring the VOC samples. Prof. Philippe Constant provided invaluable comments on early version of the manuscript.

## Author contributions

XW, EY designed the study, XW and RS performed the experiment and RS analyzed the data. RS wrote the manuscript with the revisions and comments from EY, XW and PG. All coauthors participated in discussions at the working group meetings and granted valuable advices of the manuscript.

