## Supplementary for "Nitrapyrin has far reaching effects on the soil microbial community structure, composition, diversity and functions"

**Table S1**: Reagents and reaction conditions for two PCR amplification steps for 16S and ITS sequencing.

| **Step 1: 1µl of template DNA** | | |
|---|---|---|
| **Reagent** | **Quantity (µl)** | **Reaction conditions:** |
| 10X buffer | 2.5 | 95℃ for 5.00 min;<br>30 cycles at 95℃ for 30s, 55℃ for 30s, 72℃ for 1min;<br>72℃ for 5 min;<br>4℃ forever. |
| $MgCl_2$ (25mM) | 0.5 | |
| 515F (10uM) / ITS1F (10uM) | 0.5 | |
| 806R (10uM) / 58A2R (10uM) | 0.5 | |
| dNTP (10 mM) | 0.5 | |
| Taq (5U/µl) | 0.125 | |
| BSA (50 mg/ml) | 0.25 | |
| Filter Nuclease-Free $H_2O$ | Fill to final 25 | |
| **Step 2: 5µl of DNA product from step 1** | | |
| **Reagent** | **Quantity (µl)** | **Reaction conditions:** |
| 10X buffer | 2.5 | 95℃ for 5.00 min;<br>8 cycles at 95℃ for 30s, 55℃ for 30s, 68℃ for 30s;<br>68℃ for 5 min;<br>4℃ forever. |
| $MgCl_2$ (25mM) | 0.5 | |
| Index primerF (5uM) | 2 | |
| Index primerR (5uM) | 2 | |
| dNTP (10 mM) | 0.5 | |
| Taq (5U/µl) | 0.125 | |
| BSA (50 mg/ml) | 0.25 | |
| Filter Nuclease-Free $H_2O$ | Fill to final 25 | |

**Table S2**: Reagents and reaction conditions for qPCR of selected functional genes.

| Reagent | Quantity (µl) |
|---|---|
| 1X Master Mix (HotStar iTaq DNA Polymerase, dNTPs, MgCl2, SYBR Green I dye) | 10 |
| PrimerF (500nM) | 0.6 |
| PrimerR (500nM) | 0.6 |
| DNA template (100-fold diluted) | 5 |
| Filter Nuclease-Free $H_2O$ | Fill to final 25 |
| **Gene** | **Reaction conditions:** |
| *crenamoA*<br>*amoA* | Segment 1: 98°C for 3min;<br>Segment 2: 40 cycles at 98°C for 15s, 61°C for 50s, 72°C for 1min;<br>Segment 3: 95°C for 30s, 65°C for 30s, 95°C for 30s; |
| *nirK*<br>*nirS*<br>*nifH* | Segment 1: 95°C for 1min;<br>Segment 2: 40 cycles at 95°C for 15s, 59°C for 30s, 72°C for 30s;<br>Segment 3: 95°C for 1min, 65°C for 30s, 95°C for 30s; |

**Table S3**: List of significant VOCs ($P < 0.01$) in descending order produced by rhizosphere microbial communities across sampling dates (2019-07-23, 2019-09-05).

| Compound ID | Class | RT | RI | *P*-value |
|---|---|---|---|---|
| 1377_alpha-Pinene | Terpene | 6.93 | 938.8 | 5.28E-09 |
| 386_Hexanoic acid, methyl ester | Ester | 6.75 | 927.8 | 1.57E-06 |
| 325_Unknown | | 2.04 | 613.2 | 4.18E-06 |
| 279_2,2-Diethoxyethanol | Alcohol | 2.52 | 639.4 | 5.64E-06 |
| 602_Unknown | | 7.07 | 947.3 | 1.60E-05 |
| 1099_Toluene | Alcohol | 4.34 | 769.4 | 2.66E-05 |
| 364_Isopropyl acetate | Ester | 3.13 | 680.7 | 4.59E-05 |
| 638_Unknown | | 14.57 | 1459.9 | 1.41E-04 |
| 1072_Unknown | | 12.68 | 1316.5 | 2.57E-04 |
| 483_Carbamic acid, methyl-, ethylester | Ester | 6.28 | 898.9 | 2.67E-04 |
| 107_1-Hexene, 4-methyl- | Alkene | 2.65 | 647.6 | 3.38E-04 |
| 38_Methyl acetate | Ester | 2.38 | 630.9 | 3.92E-04 |
| 1058_1,8-Octanediol | Alcohol | 9.68 | 1109.3 | 8.28E-04 |

RI: Linear retention index of 30 × 0.25 (Film 0.25) DM-5MS-UI column Agilent; RT-Retention time.

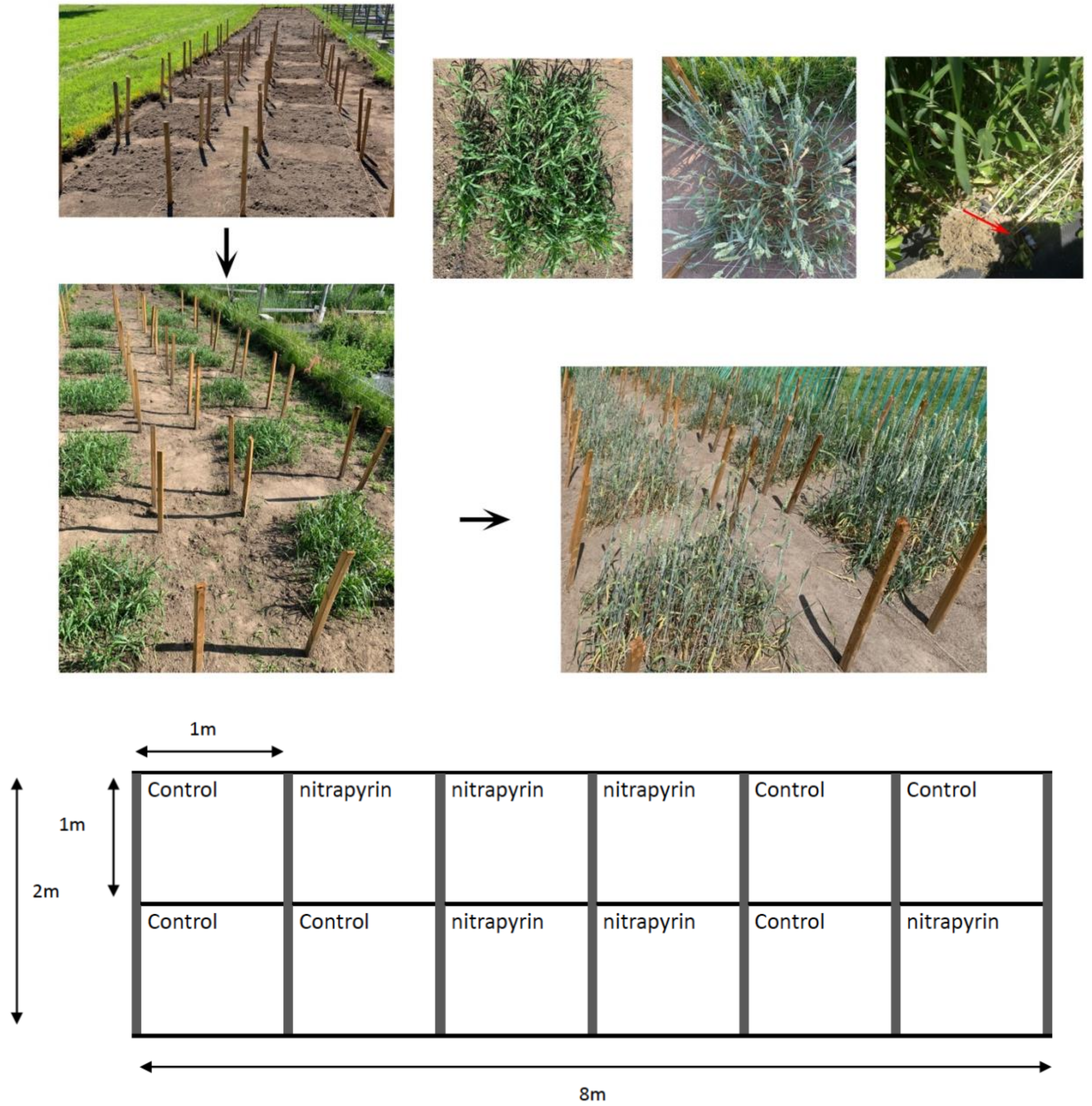


**Figure S1**: The field trial of nitrification inhibition set up in June 2019 at the Armand-Frappier Santé Biotechnologie Research Center (Laval, Québec, Canada). Control: Fertilizer without nitrapyrin. Nitrapyrin: Fertilizer with nitrapyrin. Plot size was 1m×1m, with 6 replicates per treatment. The top right picture shows the VOC sampling with PDMS tubes, stored in HPLC vials.
